# ALS iPSC-derived microglia and motor neurons respond to astrocyte-targeted IL-10 and CCL2 modulation

**DOI:** 10.1101/2023.08.14.553284

**Authors:** Reilly L. Allison, Allison D. Ebert

## Abstract

Amyotrophic lateral sclerosis (ALS) is a fatal neurodegenerative disease characterized by the loss of upper and lower motor neurons (MNs). The loss of MNs in ALS leads to muscle weakness and wasting, respiratory failure, and death often within two years of diagnosis. Glial cells in ALS show aberrant expression of pro-inflammatory and neurotoxic proteins associated with activation and have been proposed as ideal therapeutic targets. In this study, we examined astrocyte-targeted treatments to reduce glial activation and neuron pathology using cells differentiated from ALS patient-derived iPSC carrying SOD1 and C9ORF72 mutations.

Specifically, we tested the ability of increasing interleukin 10 (IL-10) and reducing C-C motif chemokine ligand 2 (CCL2 / MCP-1) signaling targeted to astrocytes to reduce activation phenotypes in both astrocytes and microglia. Overall, we found IL10/CCL2NAb treated astrocytes to support anti-inflammatory phenotypes and reduce neurotoxicity, though through different mechanisms in SOD1 and C9ORF72 cultures. We also found altered responses of microglia and motor neurons to astrocytic influences when cells were cultured together rather than in isolation. Together these data support IL-10 and CCL2 as non-mutation-specific therapeutic targets for ALS and highlight the role of glial-mediated pathology in this disease.

**Main Points (2-3 bulleted sentences, 250 characters including spaces):**

- Astrocyte-targeted IL-10 and neutralization of CCL2 reduce neuroinflammatory phenotypes in SOD1 and C9ORF72 models of ALS through divergent pathways.
- ALS microglia and motor neurons in purified monocultures display altered reactivity to pro– and anti-inflammatory influences from astrocytes compared to co-cultures.

## Introduction

Amyotrophic lateral sclerosis (ALS) is a fatal neurodegenerative disease. The primary phenotype of ALS is progressive loss of both upper and lower motor neurons resulting in muscle weakness, paralysis, and death within 2-5 years of diagnosis^1^. Though the vast majority of ALS cases are sporadic in nature with no genetic cause, there are inherited cases of familial ALS. Mutations in over 40 causative genes have been identified in familial ALS, the most common of which are mutations in superoxide dismutase 1 (SOD1) and chromosome 9 open reading frame 72 (C9ORF72)^2^. The mechanisms of how these mutations contribute to disease progression and lead to motor neuron degeneration are still unclear. Previous studies have identified the ability of both astrocytes and microglia to contribute to motor neuron toxicity in ALS^3,4^, and glial activation can be localized to degenerating motor neurons (MNs) and associated with disease progression in ALS patients^5^. ALS patient-derived induced pluripotent stem cells (iPSCs) differentiated into astrocytes, microglia, and MNs have allowed for characterization of mutation-specific ALS phenotypes. Clear differences have been identified in the transcriptional profiles of iPSC-differentiated MNs carrying SOD1 and C9ORF72 mutations^6^, setting a precedence for mutation-specific mechanisms of pathology in ALS.

These mutation-specific phenotypes may clarify contrasting theories regarding whether glial pathology is initiated by astrocytes or microglia in ALS. In support of astrocyte-driven pathology, MNs exposed to C9ORF72 astrocytes have decreased firing^7^, increased oxidative stress^4^, and increased cell death^4,8^. There were early reports of SOD1 astrocyte contribution to MN loss^3^ and improvements to lifespan when mutant SOD1 was knocked down in astrocytes^9^; however, these data were from *in vivo* studies where microglia are also present. Interestingly, SOD1 microglial activation was found to occur before disease onset in SOD1 mice^10^, and targeting these pro-inflammatory microglia^10,11^ or their production of astrocyte-activating factors^12^ has shown to extend survival. A recent transcriptomic study comparing across human and mouse models of ALS attributed production of reactive oxygen species and overall senescent phenotypes to C9ORF72 astrocytes, whereas SOD1 astrocytes were found to upregulate pathways involved in reactivity to stimuli^13^. These data support differing astrocyte-driven dysfunction in C9ORF72 models while microglia appear to initiate aberrant activation in SOD1 models.

Increased inflammation is shared characteristic across ALS mouse models and patient samples^14^, and C-C motif chemokine ligand 2 (CCL2/ MCP-1) has been specifically identified in patient CSF^15–18^, and its expression localized to astrocytes in patient spinal cord^19^. Importantly, this CCL2 upregulation is found in both familial and sporadic ALS patients; therefore, it may be an ideal candidate for therapeutics. We have previously found microglia secreting high levels of the anti-inflammatory cytokine interleukin 10 (IL-10) capable of reducing astrocyte-induced ALS phenotypes in a sporadic ALS patient iPSC model^20^. Anti-inflammatory microglia expressing high levels of IL-10 were also found to delay disease onset and extend survival in an SOD1 mouse model^21^. In this study, we differentiated astrocytes, microglia, and motor neurons from ALS patient-derived iPSCs carrying SOD1 and C9ORF72 mutations in order to compare the effects of astrocyte-targeted anti-inflammatory treatments. Specifically, we hypothesized that increasing IL-10 and reducing CCL2 signaling in astrocytes would reduce glial-driven inflammation and motor neuron dysfunction in both SOD1 and C9ORF72 cultures. IL-10 treated astrocytes have shown to reduce microglial activation better than treating microglia directly^22^, so this treatment paradigm was designed to address both microglial and astrocyte dysfunction. We confirmed the overexpression of CCL2 by C9ORF72 astrocytes and found that SOD1 astrocytes express CCL2 after exposure to SOD1 microglia. IL10/CCL2NAb treatment induced less reactive phenotypes for both C9ORF72 and SOD1 conditions, although specific signaling pathways altered by the treatment and the downstream impact on microglia and MNs diverged between SOD1 and C9ORF72 backgrounds. These data reveal the importance of paracrine signaling between glial cells and neurons in ALS and support the potential for astrocyte-targeted anti-inflammatory treatments to reduce ALS pathology.

## Results

Spinal cord patterned astrocytes were differentiated using established protocols^20,23,24^ from ALS patient-derived induced pluripotent stem cells (iPSCs) diagnosed with familial ALS due to a mutation of superoxide dismutase 1 (*SOD1*, A4V) or *C9ORF72* genes. SOD1 and C9ORF72 astrocytes appeared morphologically similar (Figure 1A) and expressed consistent transcript levels for astrocyte markers including glial fibrillary acidic protein (GFAP) and aldehyde dehydrogenase 1 family member L1 (ALDH1L1, Figure 1B). There were no differences in transcript expression for pro-inflammatory mediator NFkB, ligands interleukin 1 beta (IL-1β) and interleukin 6 (IL-6), or complement component C3 and C1q (Figure 1C-E, 2-way ANOVAs, ns). Transcripts for glial-derived and brain-derived neurotrophic factors (GDNF, BDNF) were also similarly expressed in baseline cultures of SOD1 and C9ORF72 astrocytes (Figure 1F, 2-way ANOVA, ns).

**Figure 1.**
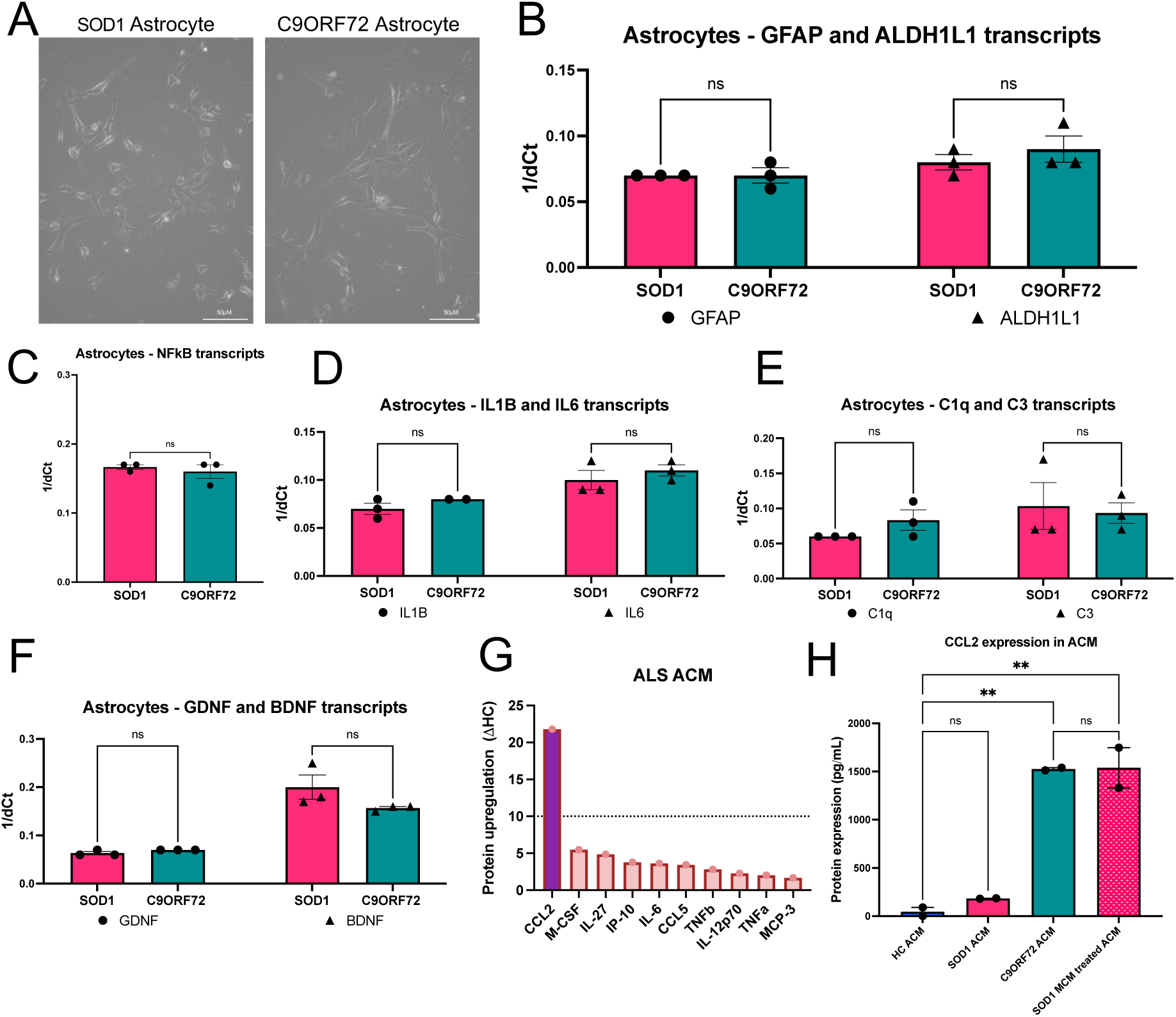
– iPSC-differentiated astrocytes from ALS patients with SOD1 and C9ORF72 mutations have similar inflammatory phenotypes. **A**. ALS patient-derived iPSCs carrying either SOD1 (left) or C9ORF72 (right) mutations differentiated into astrocytes have similar morphology (representative image, 20x objective) and expression of astrocyte marker transcripts GFAP and ALDH1L1 (**B**, 2-way ANOVA, ns). **C**. Transcripts for NFkB and downstream pro-inflammatory ligands IL-1B and IL-6 (**D**) are readily detected in both SOD1 and C9ORF72 astrocytes with no differences in expression (2-way ANOVAs, ns). **E**. Transcripts for complement components C1q and C3 as well as neurotrophic factors (**F**, GDNF, BNDF) have no differences in expression between SOD1 and C9ORF72 astrocytes (2-way ANOVA, ns). **G**. Grouped ALS astrocyte conditioned media (ACM) was found to contain 20-fold higher expression of pro-inflammatory ligand CCL2 compared to healthy control (HC) ACM. **H**. Though SOD1 ACM had similar expression of CCL2 to HCs at baseline, increased secretion of CCL2 was induced to the level of C9ORF72 ACM when SOD1 astrocytes were exposed to media conditioned by SOD1 iPSC-derived microglia (1-way ANOVA, **p<0.005).

To explore the secretome profiles of ALS astrocytes, we examined astrocyte conditioned media (ACM) via multiplex human cytokine array. Baseline (untreated) SOD1 and C9ORF72 ACM were pooled with ACM from a sporadic ALS line and compared to healthy control (HC) ACM to assess top dysregulated cytokines based on disease status regardless of mutation. We identified C-C motif chemokine 2 (CCL2, also known as MCP-1) as the top dysregulated cytokine with approximately 20-fold increased expression in ALS ACM compared to HC ACM (Figure 1G). Analysis of CCL2 expression by mutation revealed that SOD1 astrocytes had similar levels of expression to HC astrocytes at baseline. However, SOD1 astrocytes could be induced to expression levels observed in baseline C9ORF72 astrocytes when SOD1 astrocytes were exposed to media conditioned by SOD1 microglia (Figure 1H, 1-way ANOVA, HC vs SOD1 p=0.8004, HC vs C9ORF72 **p=0.0021, HC vs SOD1 MCM treated **p=0.0020). Importantly, these data are consistent with previous reports of high CCL2 expression in ALS patients^15–18^.

Because of the proposed role of microglia in ALS pathology^10^, we next investigated the influence of ACM on activation phenotypes and phagocytic ability of SOD1 and C9ORF72 iPSC-differentiated microglia (Figure 2A). Microglia were exposed to 100ng/mL lipopolysaccharide (LPS), HC ACM, mutation-specific ALS ACM (SOD1 ACM or C9ORF72 ACM), or no treatment (UTX) for 24 hours and assessed for changes in soma size as an indication of priming^25–27^. SOD1 microglial soma size increased in response to pro-inflammatory LPS and decreased in response to HC ACM showing a possible benefit of healthy astrocyte-secreted factors on SOD1 microglial priming (Figure 2B, 1-way ANOVA, UTX vs LPS ****p<0.0001, UTX vs HC ACM ****p<0.0001). SOD1 ACM did not reduce microglial soma size (HC ACM vs SOD1 ACM, **p=0.0011). C9ORF72 microglia were also reactive to pro-inflammatory LPS (Figure 2C, 1-way ANOVA, UTX vs LPS *p=0.0213), but they showed a decreased soma size in response to both HC and C9ORF72 ACM (UTX vs HC **p=0.0023, UTX vs C9ORF72 ACM **p=0.0019). To examine the effects of ACM on microglial phagocytosis, live imaging of SOD1 and C9ORF72 microglia was performed to quantify phagocytosis of pH-indicator beads (pHrodo) for 24 hours after treatment (Figure 2D). In SOD1 microglia, LPS induced a trend for higher phagocytosis (Figure 2E, 1-way ANOVA, UTX vs LPS p=0.0815), which was significantly higher than both HC and SOD1 ACM treatment conditions (LPS vs HC ACM**p=0.0050, LPS vs SOD1 ACM *p=0.0288). However, neither HC nor SOD1 ACM treatments significantly altered phagocytosis compared to untreated SOD1 microglia (UTX vs HC p=0.2563, UTX vs SOD1 p=0.6418). Despite effects of LPS and ACM on C9ORF72 microglial soma size, LPS, HC ACM, and C9ORF72 ACM treatments did not have any effects on C9ORF72 microglial phagocytosis (Figure 2F, 1-way ANOVA, ns).

**Figure 2.**
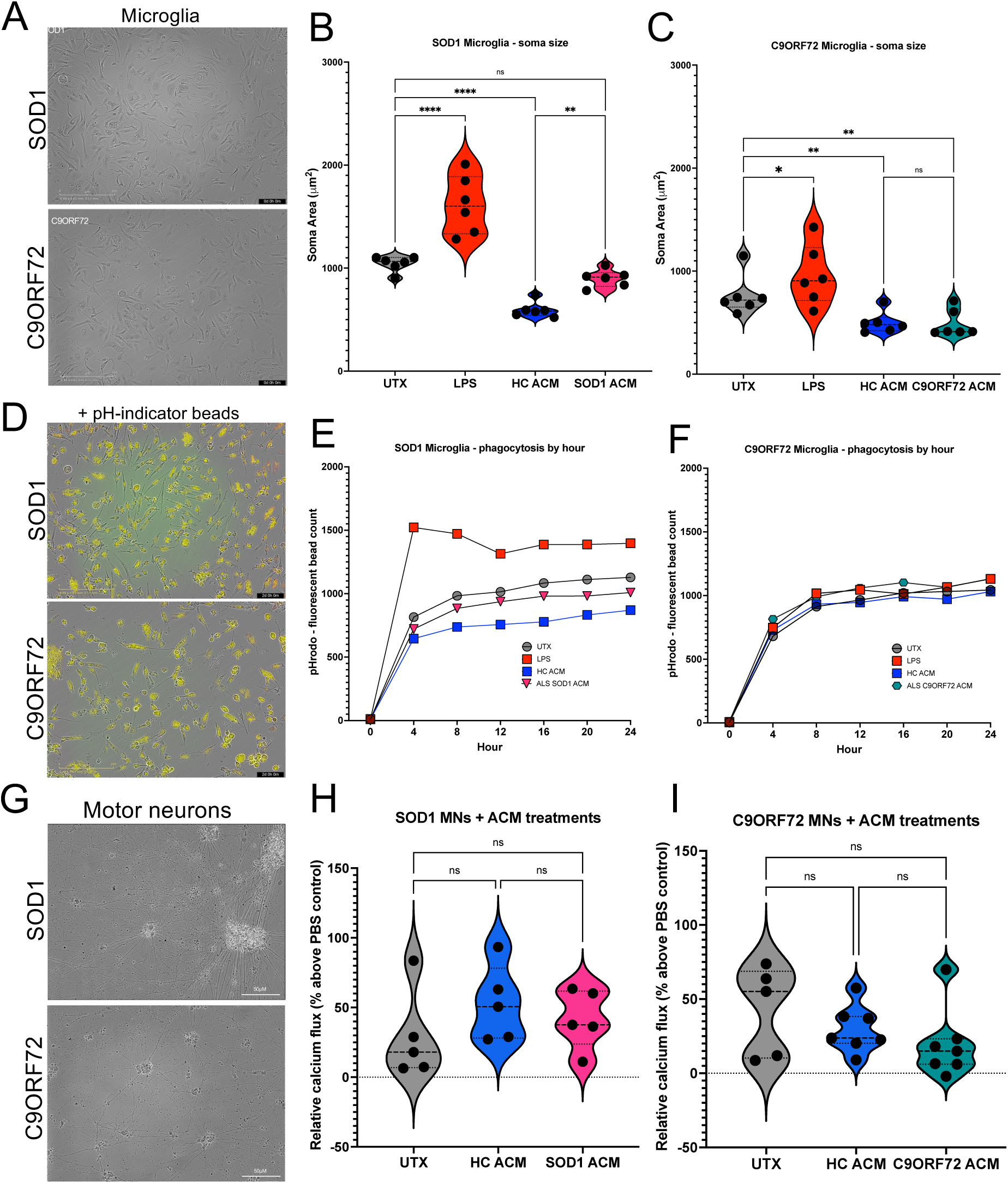
– SOD1 and C9ORF72 microglial activation and motor neuron function are not improved by exposure to ACM. SOD1 and C9ORF72 iPSC-differentiated microglia (**A**, representative image, 40x objective) have altered responses to ACM. **B**. SOD1 microglia increase soma size when treated with pro-inflammatory lipopolysaccharide (LPS) and decrease soma size when exposed to HC ACM whereas SOD1 ACM does not induce this effect (1-way ANOVA, **p<0.005, ****p<0.0001). **C**. C9ORF72 microglia increase soma size in response to LPS and decrease soma size in response to both HC and C9ORF72 ACM (1-way ANOVA, *p<0.05, **p<0.005). **D.** Phagocytosis of pH-indicator beads (pHrodo, 40x objective) is significantly decreased in SOD1 microglia by both HC (**p<0.005) and SOD1 ACM (*p<0.05) compared to LPS-treated but not to untreated SOD1 microglia (**E**, 1-way ANOVA, ns). **F**. C9ORF72 microglial phagocytosis was not influenced by exposure to LPS, HC ACM, or C9ORF72 ACM (1-way ANOVA, ns). **G**. Neither SOD1 or C9ORF72 iPSC-derived motor neurons (representative images, 20x objective) increased calcium flux in response to depolarization with KCl after exposure to HC or ALS ACM (**H**, **I,** 1-way ANOVAs, ns).

SOD1 and C9ORF72 iPSCs were next differentiated in motor neurons (MNs) to determine the influence of astrocyte secreted factors on MN function (Figure 2G). Several groups have shown iPSC-derived SOD1 and C9ORF72 MNs have increased cell death^28–30^ and abnormal function^28,31,32^ compared to control iPSC-derived MNs. To assess the direct effects of astrocyte-secreted factors on these phenotypes, MNs were treated with HC ACM, ALS ACM (SOD1 or C9ORF72 ACM), or no treatment (UTX). After 48 hours of treatment, SOD1 and C9ORF72 MNs were assessed for calcium flux in response to depolarization with KCl. In these MN monocultures, neither HC nor disease ACM was able to significantly alter the calcium response of SOD1 or C9ORF72 MNs compared to untreated cultures (Figures 2H, 2I, 1-way ANOVAs, ns). There was a trend for increased calcium flux in SOD1 MNs with HC ACM compared to UTX (p=0.1524), but this was not found with SOD1 ACM treatment.

Based on the recent finding that microglial activation is better reduced by anti-inflammatory astrocytes than direct microglial treatments^22^, we proposed that astrocyte-directed treatments would appropriately reduce both astrocyte and microglial-driven pathology in ALS. To test this hypothesis, we treated SOD1 and C9ORF72 astrocytes with 430pg/mL of recombinant human interleukin 10 (IL-10). IL-10 is shown to increase anti-inflammatory phenotypes through inhibition of NFkB and by NFkB-independent mechanisms^33–35^ as well as provide direct trophic support to neurons and reduce microglial pro-inflammatory cytokine production^36,37^. To ensure reduction of microglial activation, we combined this IL-10 treatment with 2ng/mL of CCL2 neutralizing antibodies (NAb) (IL10/CCL2NAb, Figure 3A). A 48-hour wash condition was also included to control for IL-10 or CCL2 NAb remaining in ACM and to assess non-continuous treatment on astrocyte phenotypes. BDNF transcripts were increased in C9ORF72 IL10/CCL2NAb-treated astrocytes (Figure 3B, 2-way ANOVA, UTX vs IL10/CCL2NAb *p=0.0285) and a trend toward increase was also seen in treated SOD1 astrocytes (UTX vs IL10/CCL2NAb p=0.1036). IL10/CCL2NAb C9ORF72 astrocytes also had decreased C1q transcripts that remained decreased 48-hours after treatment (Figure 3C, 2-way ANOVA, UTX vs IL10/CCL2NAb *p=0.0208, UTX vs 48wash *p=0.0420), whereas SOD1 astrocytes had no changes to C1q transcript production (ns). A decrease (SOD1, *p=0.0495) or trend for decrease (C9ORF72, p=0.0602) in IL-1β transcripts was seen in all treated astrocytes 48-hours after IL10/CCL2NAb treatment (Figure 3D, 2-way ANOVA). Interestingly, IL10/CCL2NAb C9ORF72 astrocytes increased transcripts for IL-6, which could be either neuroprotective or pro-inflammatory in ALS^38^, though this effect was not found in the 48wash condition (Figure 3E, 2-way ANOVA, UTX vs IL10/CCL2NAb **p=0.0015, UTX vs 48wash p=0.2698) nor in treated SOD1 astrocytes (ns).

**Figure 3.**
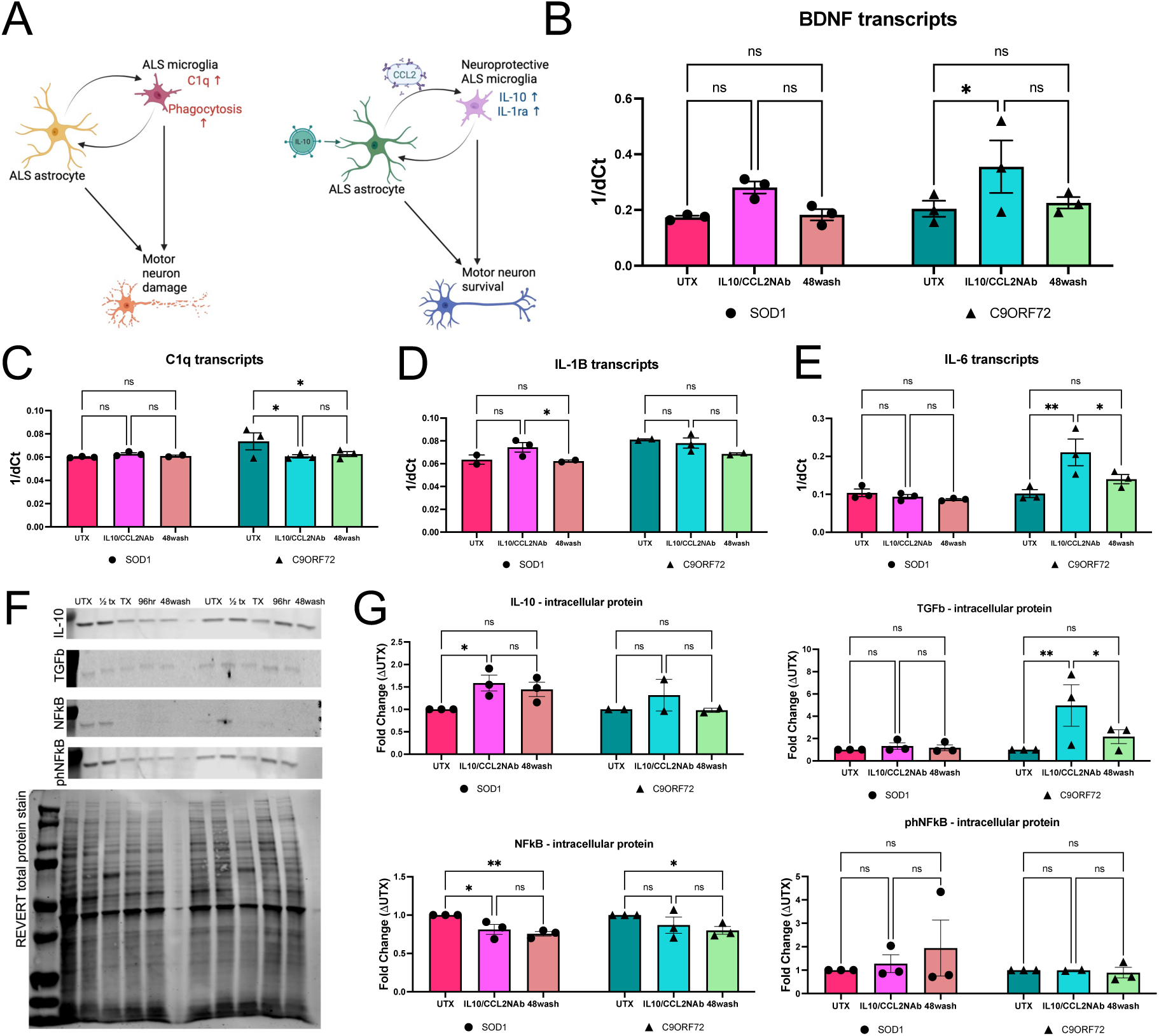
– Increase of IL-10 and neutralization of CCL2 differently but beneficially alter SOD1 and C9ORF72 astrocyte phenotypes. **A**. Schematic of ALS glial-mediated activation and motor neuron damage (left) and hypothesized effects of astrocyte-targeted treatments with 430pg/mL recombinant human IL-10 and 2ng/mL of neutralizing CCL2 antibodies (right, IL10/CCL2NAb). **B**. C9ORF72 astrocytes increased expression of BDNF transcripts with IL10/CCL2NAb and decreased transcripts for C1q (**C**) both during treatment and 48 hours after treatment was removed (48wash), while SOD1 astrocytes had no changes in expression of these targets (2-way ANOVAs, *p<0.05, ns). **D**. SOD1 astrocytes did show decreased transcript expression for IL-1β 48 hours after IL10/CCL2NAb (2-way ANOVA, *p<0.05) **E**. C9ORF72 astrocytes also had a surprising increase in IL-6 expression during IL10/CCL2NAb, though this effect was ameliorated in 48wash condition (2-way ANOVA, **p<0.005, *p<0.05). **F**. Western blot analyses identify increased anti-inflammatory and decreased pro-inflammatory protein expression after IL10/CCL2NAb in SOD1 and C9ORF72 astrocytes. IL10/CCL2NAb treatment increased protein expression of IL-10 in SOD1 astrocytes and TGFb in C9ORF72 astrocytes (**G**, 2-way ANOVA, *p<0.05, **p<0.005). Both SOD1 and C9ORF72 astrocytes had decreased protein expression of pro-inflammatory mediator NFkB 48 hours after IL10/CCL2NAb treatment, though no changes to phosphorylated NFkB (phNFkB, 2-way ANOVA, **p<0.005, *p<0.05, ns).

We next evaluated intracellular changes in anti– and pro-inflammatory protein expression via Western blot on collected cell pellets of IL10/CCL2NAb treated astrocytes (Figure 3F). Production of IL-10 by SOD1 astrocytes was increased with IL10/CCL2NAb treatment and had a trend for sustained increased in 48wash condition (Figure 3G, 2-way ANOVA, UTX vs IL10/CCL2NAb *p=0.0207, UTX vs 48wash p=0.0624). C9ORF72 astrocytes did not significantly increase IL-10 protein production, but treatment did increase anti-inflammatory TGF-β protein (Figure 3G, 2-way ANOVA, UTX vs IL10/CCL2NAb **p=0.0050). Both SOD1 and C9ORF72 astrocytes had decreased NFkB protein 48-hours after treatment (Figure 3G, 2-way ANOVA, SOD1 UTX vs 48wash **p=0.0096, C9ORF72 UTX vs 48wash *p=0.0274) and no changes to phosphorylated NFkB (ns). Although signaling pathways activated by IL10/CCL2NAb treatment may be mutation-specific, patterns of decreased pro-inflammatory and increased anti-inflammatory signaling are apparent in both SOD1 and C9ORF72 IL10/CCL2NAb-treated astrocytes.

To assess the efficacy of astrocyte-targeted IL10/CCL2NAb treatment on microglial activation and phagocytosis, we treated SOD1 and C9ORF72 microglia with ACM from IL10/CCL2NAb and 48wash astrocytes for 24 hours (Figure 4A). SOD1 microglia responded to SOD1 IL10/CCL2NAb ACM and 48wash ACM with a decrease in soma size compared to SOD1 ACM (Figure 4B, 1-way ANOVA, p****p<0.0001), whereas the already small C9ORF72 microglia soma size was not negatively impacted by IL10/CCL2NAb treatment (Figure 4C, 1-way ANOVA, ns). SOD1 microglia had a decreased trend for phagocytosis of pH-indicator beads with SOD1 48wash ACM (Figure 4D, 4E, 1-way ANOVA, UTX vs IL10/CCL2NAb p=0.2219, UTX vs 48wash p=0.0589). Although not quite significant, the separation of C9ORF72 microglial phagocytosis after exposure to IL10/CCL2NAb and 48wash is notable compared to the LPS and HC ACM treated groups in 2F (Figure 4E, 1-way ANOVA, UTX vs IL10/CCL2NAb ACM p=0.1458, IL10/CCL2NAb ACM vs 48wash p=0.0553).

**Figure 4.**
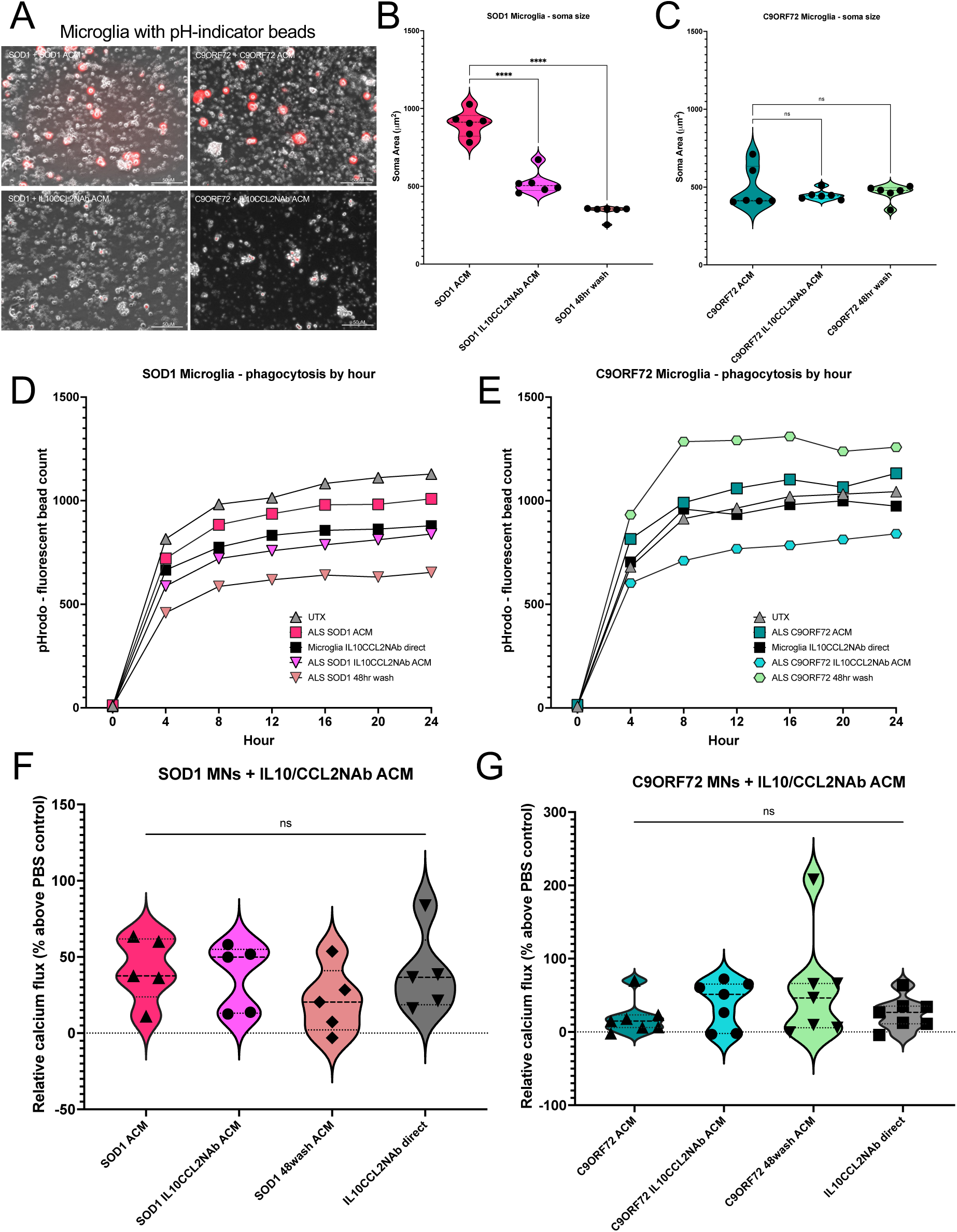
– IL10CCL2-treated astrocytes may influence microglial activation but not motor neuron function in mono-cultures. **A**. Representative image of SOD1 and C9ORF72 microglia exposed to untreated or IL10/CCL2NAb ACM (20x objective). **B**. SOD1 microglia treated with SOD1 IL10/CCL2NAb ACM do show an improvement in priming as measured by soma size (1-way ANOVA, ****p<0.0001) while C9ORF72 microglia remain in unprimed status regardless of ACM treatment (**C**, 1-way ANOVA, ns). **D**. Both IL10/CCL2NAb and 48wash ACM from SOD1 astrocytes show trends for decreased phagocytosis of pH-indicator beads by SOD1 microglia compared to untreated (1-way ANOVA, p>0.05, while only IL10/CCL2NAb ACM has a trend for decreased phagocytosis in C9ORF72 microglia (**E**, 1-way ANOVA, p>0.05). Direct treatment of microglia with IL-10 and CCL2 neutralizing antibodies did have a similar effect on SOD1 microglia as IL10/CCL2NAb ACM, while it did not change phagocytic ability of C9ORF72 microglia (1-way ANOVAs, SOD1: IL10/CCL2NAb vs direct treatment p=0.7820, C9ORF72: UTX vs direct treatment p=0.9439). Neither SOD1 (**F**) nor C9ORF72 (**G**) motor neurons had increased calcium flux after exposure to IL10/CCL2NAb ACM, 48wash ACM, or direct IL10CCL2 treatments compared to untreated ACM (1-way ANOVAs, ns). SOD1 ACM and C9ORF72 ACM values in B, C, F, and G repeated from Figure 2 for ease of comparison.

To compare astrocyte-targeted vs microglial-targeted treatments, we also applied 430pg/mL IL-10 with 2ng/mL CCL2 NAbs directly onto microglia with ALS ACM treatments. SOD1 microglia with direct IL10/CCL2NAb treatments had similar rates of fluorescent bead phagocytosis to SOD1 I IL10/CCL2NAb ACM, but only half the effect of SOD1 48wash ACM (Figure 4D mean differences: SOD1 ACM vs IL10/CCL2NAb ACM = 145 beads, SOD1 ACM vs direct IL10/CCL2NAb = 92.44 beads, SOD1 ACM vs 48wash = 210.8 beads). Direct IL10/CCL2NAb treatments on C9ORF72 microglia were about one third as effective as astrocyte-targeted treatments (Figure 4E mean differences: C9ORF72 ACM vs IL10/CCL2NAb ACM = 235.8 beads, C9ORF72 ACM vs direct IL10/CL2NAb = 88.64 beads). Application of IL10/CCL2NAb ACM, 48wash ACM, or direct IL10/CCL2NAb treatments of SOD1 or C9ORF72 MN monocultures did not increase calcium flux compared to untreated ACM (Figure 4F, 4G, 1-way ANOVAs, ns). Together, these data suggest that astrocyte-targeted IL10/CCL2NAb treatment can reduce activation of SOD1 and C9ORF72 microglia and do so more effectively than microglia-directed treatments. However, improvements to ALS MN function may require influences greater than short term exposure to astrocyte secreted factors.

To further address the impacts of glial modulation on MNs, we examined the effects of astrocyte-targeted treatments in co-cultures of MNs and microglia to assess the importance of dynamic cell interactions on therapeutic effects. To allow for live cell imaging, we established stable fluorescent lines of SOD1 and C9ORF72 iPSCs using lentiviruses driving expression of either GFP or RFP under the elongation factor 1 (EF-1) promotor. These were then separately differentiated into GFP+ or RFP+ microglia and MNs before microglia were introduced to MN cultures (Figure 5A). A trend for increased calcium flux was found in SOD1 MN + SOD1 microglia co-cultures when treated with SOD1 IL10/CCL2NAb ACM compared to untreated or SOD1 ACM treated cultures (Figure 5B, 1-way ANOVA, UTX vs IL10/CCL2NAb p=0.1006, SOD1 vs IL10/CCL2NAb p=0.1362). In C9ORF72 co-cultures, C9ORF72 ACM significantly reduced calcium flux, an effect that was ameliorated by C9ORF72 IL10/CCL2NAb ACM (Figure 5B, 1-way ANOVA, UTX vs C9ORF72 ACM *p=0.0224, UTX vs IL10/CCL2NAb ACM ns). This is likely due to astrocyte-driven microglial activation, as these effects were not seen in earlier MN monocultures treated with C9ORF72 ACM.

**Figure 5.**
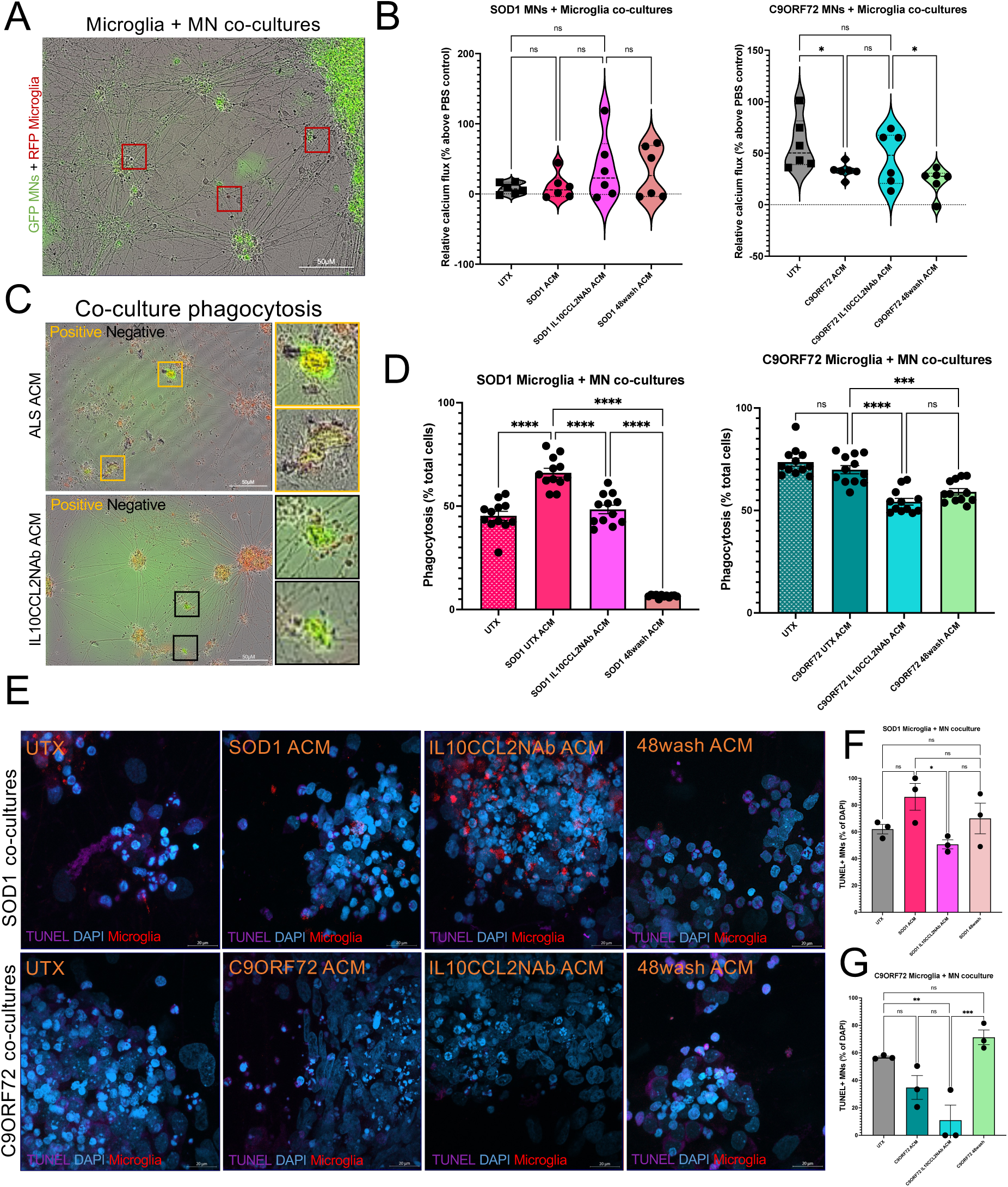
– IL10/CCL2NAb treated astrocytes reduce phagocytosis and apoptosis of SOD1 and C9ORF72 motor neurons in co-cultures with microglia. **A**. GFP– and RFP-labeled MNs and microglia allow for live-imaging analyses of co-cultures (20x objective). **B**. Exposure of SOD1 co-cultures to SOD1 ACM does not alter calcium flux, but a trend for increased flux can be found after IL10/CCL2NAb ACM exposure compared to untreated and SOD1 ACM co-cultures (UTX vs IL10/CCL2NAb p=0.1006, SOD1 ACM vs IL10/CCL2NAb p=0.1362). C9ORF72 co-cultures significantly decrease flux after C9ORF72 ACM treatment, with a trend for amelioration with IL10/CCL2NAb ACM treatment (UTX vs IL10/CCL2NAb p=0.2145, C9ORF72 ACM vs IL10/CCL2NAb p=0.2566). **C**. Phagocytosis of co-cultures quantified using RFP+/GFP+ colocalization in microglia (yellow box – GFP/RFP double positive, black box – not double positive) after exposure to ACM via live imaging paradigm. **D**. SOD1 ACM increases phagocytosis in SOD1 co-cultures which is ameliorated by IL10/CCL2NAb treatment and 48-hours afterwards (1-way ANOVA, ****p<0.0001). C9ORF72 ACM does not increase already high levels of phagocytosis observed in untreated C9ORF72 co-cultures, but this rate is significantly decreased by both IL10/CCL2NAb and 48wash ACM (1-way ANOVA, ***p<0.0005, ****p<0.0001). **E**. TUNEL assay to assess MN apoptosis in SOD1 and C9ORF72 MN + Microglia co-cultures (63x objective). **F**. IL10/CCL2NAb ACM decreases apoptosis compared to ALS ACM in both SOD1 and C9ORF72 co-cultures (1-way ANOVAs, *p<0.05, ***p<0.0001).

To test this hypothesis, we quantified phagocytosis in these co-cultures as microglia became double-positive for RFP and GFP upon engulfment of fluorescent MN elements (Figure 5C). 48 hours after co-cultures were treated with ACM, GFP+/RFP+ microglia were significantly increased in SOD1 co-cultures with SOD1 ACM and significantly reduced in both SOD1 IL10/CCL2NAb and 48wash ACM (Figure 5D, 1-way ANOVA, UTX vs SOD1 ****p<0.0001, SOD1 vs IL10/CCL2NAb ****p<0.0001, SOD1 vs 48wash ****p<0.0001). C9ORF72 co-cultures had a high rate of phagocytosis that was not increased by C9ORF72 ACM, but both C9ORF72 IL10/CCL2NAb and 48wash ACM reduced these values (Figure 5D, 1-way ANOVA, UTX vs C9ORF72 ACM p=0.4659, C9ORF72 ACM vs IL10/CCL2NAb ****p<0.0001, C9ORF72 ACM vs 48wash ***p=0.0006). Further, we found that addition of microglia without astrocyte influence did not significantly alter calcium flux in either SOD1 or C9ORF72 MNs compared to MN monocultures (Supplemental Figure S2, t-tests, ns).

Finally, we examined the effects of IL10/CCL2NAb astrocyte-targeted treatments on MN cell death in co-cultures using a TUNEL stain (Figure 5E, Supplemental Figure S3). We identified high baselines of MN cell death in untreated co-cultures with microglia in both SOD1 and C9ORF72 cultures (Figure 5F). Exposure of SOD1 co-cultures to SOD1 ACM increased TUNEL+ MNs compared to SOD1 IL10/CCL2NAb ACM (1-way ANOVA, UTX vs SOD1 ACM p=0.0732, SOD1 ACM vs IL10/CCL2NAb *p=0.0147). C9ORF72 co-cultures had reduced MN death when treated with C9ORF72 IL10/CCL2NAb ACM compared to UTX, though this effect was not sustained in 48wash ACM condition (1-way ANOVA, UTX vs IL10/CCL2NAb **p=0.0054, IL10/CCL2NAb vs 48wash ***p=0.0007). Together, these data highlight convergent and divergent effects of glial-neuron interactions in both SOD1 and C9ORF72 iPSC cultures.

## Discussion

Although only 10% of ALS patients have familial cases that can be linked to mutations in causative genes, these have historically been the focus of research^39,40^. This is primarily due to the logistics of creating animal models for sporadic ALS without a specific genetic mutation to target. However, growing evidence supports the idea that the specific mechanisms of motor neuron loss may not be shared across ALS inherited mutations and sporadic cases^6,41^. iPSC models of disease have allowed for more exploratory and comparative studies to begin addressing this. For example, SOD1 and C9ORF72 mutation carrying iPSC-differentiated motor neurons have been shown to have unique transcriptional profiles, with defects in RNA processing and transport unique to C9ORF72 MNs and SOD1 MNs downregulating TGFβ, SMAD, and MAPK signaling^6^. Nevertheless, there are commonalities across ALS backgrounds. In addition to loss of upper and lower motor neurons in ALS, one shared hallmark of disease across familial and sporadic cases is glial-mediated inflammation^3–5^. Though it may be either microglia or astrocytes that initiate this activation, it has been proposed that converting these glial cells towards a more neuroprotective phenotype can prevent disease onset and symptom progression^42–46^. Based on this concept, we hypothesized that astrocyte-targeted therapeutics would prevent both astrocyte-driven neurotoxicity and promote anti-inflammatory microglial phenotypes.

In this study, we examined IL-10 as an anti-inflammatory therapeutic targeted to astrocytes carrying SOD1 and C9ORF72 mutations. We combined IL-10 expression with a neutralizing antibody treatment against CCL2 – a cytokine capable of inducing microglial activation^47^ that we and others have found highly upregulated by ALS astrocytes^15–19^ (Figure 1). Previous studies have tried directly targeting activated microglia with IL-10 treatments and found it ineffective without other anti-inflammatory treatments^48^ or astrocyte-secreted factors^22^, so we focused on astrocyte-directed anti-inflammation. Overall, we saw reduced pro-inflammatory factors, increased anti-inflammatory signaling, and increased neurotrophic support from IL10/CCL2NAb treated astrocytes (Figures 2, 3), though the specific pathways activated were different between SOD1 and C9ORF72 cultures. At baseline, C9ORF72 astrocytes appeared to reside in a more pro-inflammatory state than SOD1 astrocytes, whereas SOD1 microglial were more reactive to astrocytic influences than C9ORF72 microglia (Figure 2). Though microglia do express C9ORF72, microglia with mutations in C9ORF72 have recently been reported to be very similar to healthy controls unless exposed to extrinsic factors^49^, thereby supporting an astrocyte-driven activation in C9ORF72-associated ALS. Though the exact function of the C9ORF72 is unclear, it is thought to interact with Rab proteins to regulate endosomal trafficking, autophagy, and lysosomal biogenesis^50^. Mutations in C9ORF72 in astrocytes are associated with abnormal RNA metabolism, increased NFkB, and altered glutamate regulation all of which can contribute directly to increased neurotoxicity^7,51^.

Fitting with *in vivo* datasets of upregulated CCL2 by SOD1 astrocytes^19^, we found that SOD1 astrocytes could be induced to a pro-inflammatory status after exposure to SOD1 microglia conditioned media (Figure 1H). A recent study found SOD1 astrocytes more reactive to pro-inflammatory stimuli than other ALS astrocytes^52^, which supports the idea of microglia-driven activation in SOD1-associated ALS. This microglial activation could result from both gain and loss of function mechanisms due to SOD1 protein, as SOD1 is an antioxidant enzyme responsible for regulation of reactive oxygen species^53^ (ROS). Microglia monitor ROS^54^ and increase Toll-like receptor (TLR)-mediated inflammatory responses in response to insufficient degradation of ROS^55,56^. This mechanism, coupled with the gain of protein aggregates that microglia are tasked with clearing^57^, may explain why SOD1 mutations are associated with activated microglia over activated astrocytes.

After IL10/CCL2NAb treatment (Figure 3), SOD1 astrocytes decreased NFkB and increased production of IL-10 – an ideal combination to reduce microglial-mediated inflammation. In support of this, application of IL10/CCL2NAb treated SOD1 ACM onto SOD1 microglia did reduce activation phenotypes (Figure 4). Treated C9ORF72 astrocytes increased TGFβ production, which early in ALS disease course is found to increase neuroprotection and decrease excitotoxicity^58^. Combined with the increased BDNF and reduced tagging of synapses for microglial engulfment through decreased C1q, IL10/CCL2NAb induced C9ORF72 phenotype appears calibrated to specifically target astrocyte-driven neurotoxicity. Expectedly, treated C9ORF72 ACM again had little impact on the homeostatic status of C9ORF72 microglia (Figure 4). We also confirmed the importance of dynamic signaling between cells on activation and function in disease contexts through the use of co-cultures (Figure 5). Although we found no effect of ACM on MN calcium flux in either SOD1 or C9ORF72 monocultures, co-cultures of motor neurons and microglia had altered calcium flux compared to MNs cultured alone.

A change in MN calcium flux can be partially attributed to microglial activation and phagocytosis, particularly in untreated SOD1 co-cultures where MN response to depolarizing stimuli is reduced compared to SOD1 MNs alone (Figure 5B vs 2H). However, in C9ORF72 co-cultures which have a high rate of phagocytosis with or without ACM treatment (Figure 5D), calcium flux is significantly decreased in co-cultures treated with C9ORF72 ACM compared to untreated co-cultures (Figure 5B). This indicates the detrimental effects of C9ORF72 astrocytes towards MNs are likely not prevented by C9ORF72 microglia, and that C9ORF72 microglia are likely more activated in the presence of MNs than in monocultures. In support of this, we found that both untreated and ACM-treated C9ORF72 co-cultures had high levels of neuron death that could be reduced by astrocyte-targeted IL10/CCL2NAb treatment. In SOD1 co-cultures, a strong trend for increased neuron death was associated with increased phagocytosis but not with decreased neuron calcium flux. This finding could indicate an increase in excitability of surviving MN populations^59,60^ as a compensatory mechanism for those lost due to microglial engulfment or cytotoxicity.

These co-culture experiments highlight the importance of dynamic communication between glial cells and neurons in the context of ALS. iPSC-differentiated monocultures remain a valuable tool to identify mechanisms of intrinsic dysfunction in diseased cells. However, therapeutics designed to correct dysfunction should be examined in both mono– and co-cultures to assess how paracrine interactions with other cell types may influence phenotypes and pathology. Overall, this *in vitro* study provides insight into the mechanisms of glial-driven pathology in SOD1 and C9ORF72 associated models of ALS. Importantly, astrocyte-targeted IL-10 and reduction of CCL2 was found beneficial in both models, indicating a potential for a broad therapeutic effect. In considering translation to *in vivo* applications, the *in vitro* 48wash data support a sustained treatment paradigm, such as through gene therapy-based approaches, as removal of treatment allowed the return of some neuroinflammatory and neurotoxic phenotypes. Together, these data support IL-10 and CCL2 as therapeutic targets for ALS patients across mutations.

## Materials and methods

### Cell culture

Two healthy control (21.8, 4.2) and three ALS patient (ALS 71 [SOD1 A4V mutation], CS29i [C9ORF72 HRE mutation], AB34.12 [sporadic ALS]) iPSC lines were utilized in these experiments^20,61–63^. All pluripotent stem cells were maintained on Matrigel (Corning) in Essential 8 (Gibco) and passaged every 4-6 days. iPSCs and differentiated cells were confirmed mycoplasma negative. SOD1 and C9ORF72 lines were made into GFP– and RFP-expressing stable lines by infecting cells with LentiBrite GFP Control Lentiviral Biosensor (Millipore, #17-10387, titer 7.34 x108 IFU/mL) or LentiBrite RFP Control Lentiviral Biosensor (Millipore, #17-10409, titer 4.59 x 108 IFU/mL) at MOI of 20 for 24 hours. Virus was removed and cells were allowed to expand for 1 week. GFP+ and RFP+ cells were isolated using WOLF Cell Sorter (Nanocellect) and expanded to create purified stable lines.

### Astrocyte, microglia, and motor neuron differentiations

Spinal cord patterned astrocytes were generated from iPSCs as previously described^20,23,24^. Differentiation reagents were purchased from ThermoFisher unless otherwise noted. Briefly, iPSCs were differentiated into neural progenitor cells (NPCs) using dual SMAD inhibition (SB 431542 and LDN 1931899) and patterned towards ventral-caudal spinal cord using retinoic acid (RA) and hedgehog smoothened agonist (SAG). NPCs were passaged every 6 days and on passage 3 were differentiated into astrocytes using ScienCell Astrocyte Medium (ScienCell Research Laboratories, Carlsbad, CA, USA) containing 1% astrocyte growth supplement, 1% penicillin–streptomycin, and 2% B27. After passage 4, astrocytes were used for ACM generation and collection.

Microglia were differentiated using the commercially available differentiation kit (StemCell Technologies #05310, #100-0019, #100-0020, Vancouver, BC, Canada) based on a previously published protocol^64^. As previously described^20,23^, iPSCs were differentiated into hematopoietic progenitor cells (HPCs) using the STEMdiff Hematopoietic Kit (StemCell Technologies, Vancouver, BC, Canada). Floating HPCs were collected and plated at 50,000 cells/mL in STEMdiff microglia differentiation media (StemCell Technologies, Vancouver, BC, Canada) for 24 days, followed by rapid maturation in STEMdiff microglia maturation media (Stem-Cell Technologies, Vancouver, BC, Canada) for a minimum of 4 days.

Spinal motor neurons were differentiated based on the Maury et al. (2015) protocol^65^. Briefly, embryoid bodies were generated from iPSCs and patterned in the presence of Chir-99021 with dual SMAD inhibition (SB 431542 and LDN 1931899) followed by treatment with retinoic acid (RA), smoothened agonist (SAG), and DAPT. Spinal motor neuron progenitor cells were then dissociated and plated on Matrigel-coated glass coverslips or 96-well plates for terminal differentiation and maturation in growth factor supplemented medium for 21–42 days *in vitro*.

### Quantitative real-time polymerase chain reaction (qRT-PCR)

RNA was isolated from cell pellets using the RNeasy Mini Kit (Qiagen) following manufacturer’s instructions, quantified using a Nanodrop Spectrophotometer, treated with RQ1 Rnase-free Dnase (Promega), and converted to cDNA using the Promega Reverse Transcription system (Promega). SYBR green RT-qPCR was performed in triplicate using cDNA and run on the Bio-Rad CFX384 real time thermocycler. Primer sequences shown in Table S1. Cq values for each target were normalized to GAPDH and calculated using the 1/dCt method. A minimum of three differentiations for each line were collected and run in technical triplicates. Individual data points represent the average of technical triplicates for each experiment.

### Multiplex human cytokine array on conditioned media

Eve Technologies (Calgary, AB, Canada) performed the 48 multiplex cytokine array assay from duplicate differentiations using conditioned medium samples generated from iPSC-derived astrocytes. Data were analyzed for fold change difference of each cytokine expression level in ALS ACM compared to HC ACM.

### Microglia soma size and pHrodo phagocytosis assays

As previously described^66^, microglia were treated with microglia maturation media (UTX), 1:2 ACM, or 100ng/mL lipopolysaccharide (LPS, Sigma Aldrich, L2018) and placed in Incucyte (Sartorius) to allow for live cell imaging with 20x objective during 24-hour treatment. Incucyte software was used to calculate average soma area for all cells in one well (5,000 microglia per well of 24-well plate) at 24-hour timepoint for soma size analyses. Points in soma size graphs represent the average of 3 technical well replicates. After 24-hour ACM treatment and soma measurement, 1ug/mL pHrodo Red Zymosan Bioparticles (ThermoFisher, #P35364) were added to microglia cultures. Plates were returned to Incucyte and imaged for 24 hours using brightfield and red fluorescent channels at 20x. Images were analyzed using Incucyte software for total number of RFP+ microglia in each well and averaged across three technical replicates. Data points on graph represent mean and standard error of the mean for experimental replicates.

### Fluo-4NW calcium flux assay

Calcium flux was measured in MNs seeded in a 96 well plate using the Fluo-4 NW Calcium Assay Kit (ThermoFisher, #F36206) per the manufacturer’s instructions. Growth medium was removed, and 100μL of dye loading solution was added to each well and incubated for 30 minutes at 37C followed by 30 minutes at room temperature. Just before measuring fluorescence, 25μL of the 50mM KCl agonist or PBS (control) were spiked into each well. Fluorescence (excitation 494nm, emission 516nm) was immediately measured using a GloMax microplate reader. As previously described^66^, fluorescence (% above PBS baseline) was calculated by subtracting the fluorescence of PBS-stimulated wells from each test well then dividing by PBS-stimulated well value and multiplying by 100. Individual data points in relative fluorescence graphs represent the average of three technical replicates.

### *In vitro* astrocyte treatments

Treatments of SOD1 and C9ORF72 astrocytes were performed with 430pg/mL recombinant IL-10 (PeproTech, #200-10) and 2ng/mL CCL2 neutralizing antibodies (Bio-Techne, #AF-279-NA) in supplemented Astrocyte media for 48 hours before collection of treatment media and cell pellets. For 48 wash conditions, astrocytes were rinsed with PBS and fed with supplemented Astrocyte media. Media were collected from 48wash astrocytes after 48 hours.

Treatment values were set based on doubling the amount of IL-10 secreted by microglia previously found capable of reducing astrocyte-driven pathology^20^ and doubling the amount of CCL2 secreted by ALS astrocytes as determined by multiplex cytokine array (Figure 1H).

### Western blot

Cell pellets were lysed by sonication with Triton X-100 and protein concentration was determined using a BCA assay (ThermoFisher). Equal amounts of protein were loaded onto 10% or 12% pre-cast Tris-HCl Mini-PROTEAN gels (Bio-Rad, Hercules, CA, USA) and proteins separated by electrophoresis, then transferred to PVDF membranes (Bio-Rad). Membranes were blocked for 1 h in Odyssey TBS blocking buffer (LI-COR), followed by overnight primary antibody incubation and 30 min secondary antibody incubation. Quantification was performed with FIJI (ImageJ, National Institutes of Health, Bethesda, MD, USA) and normalized to REVERT total protein stain (LI-COR). Primary antibodies used were IL-10 (abcam, ab133575), mouse anti-TGFβ (ThermoFisher, #MAB-16949), mouse anti-NFkB p65 (Cell Signaling, #6956) and rabbit anti-phNFkB (Cell Signaling, #3033) all at 1:1000 dilutions. Secondary antibodies used were anti-rabbit IRDye 800CW (LI-COR, 1:5000 dilution, Lincoln, NE, USA) and anti-mouse IRDye 680RD (LI-COR, 1:5000 dilution, Lincoln, NE, USA).

### Microglia and MN co-cultures

After a minimum of 4 days in STEMdiff maturation media, SOD1 and C9ORF72 microglia were added at a ratio of 1:4 to SOD1 or C9ORF72 MNs (between 21-36 days of MN maturation) with either 25% ALS UTX, IL10/CCL2NAb, or 48wash ACM from appropriate astrocyte line, direct IL10/CCL2NAb treatment with 25% ALS UTX ACM, or 25% fresh Astrocyte media (UTX). Treated co-cultures were placed in Incucyte for 48 hours and imaged every 2 hours in brightfield, RFP, and GFP channels at 20x. After 48 hours, co-cultures were fixed for TUNEL assay or analyzed for calcium flux using Fluo4-NW assay. Phagocytosis was calculated using Incucyte software to identify the number of GFP+/RFP+ double positive microglia at the 48-hour timepoint. Data points on co-culture phagocytosis graph represent the average of three technical replicates for double positive GFP+/RFP+ microglia normalized to total number of cells in the field of view.

### TUNEL assay

Plated cells were fixed in 4% PFA for 20 min at room temperature, rinsed with PBS, and then stained using a Click-iT TUNEL Alexa Fluor 647 Assay Kit (ThermoFisher, #C10247) following manufacturer’s instructions. As previously described^66^, cells were permeabilized as described for ICC, washed, and incubated with DNA labeling solution for 1 hour at 37C. Cells were washed and optional DAPI nuclear counterstain was then applied for 30 min at room temperature. Coverslips were imaged with standard fluorescent microscopy. Three images were acquired from randomly selected fields for each coverslip. Images were analyzed for total fluorescence in either channel using FIJI (ImageJ) software. Relative expression for each condition (total TUNEL stain divided by total 7AAD stain) was analyzed to account for variable number of cells in each ROI as previously described^23^. Representative images were acquired on a Zeiss confocal microscope using 63x oil objective and are displayed as a maximum intensity projection of z-stack image series.

### Statistical analyses

Experimental conditions within each experiment were performed in technical triplicates for a minimum of three independent experiments unless otherwise noted. Data were analyzed using GraphPad Prism software and the appropriate statistical tests including the Student’s t test, 1-way ANOVA, and 2-way ANOVA followed by Tukey’s post hoc analysis of significance. Changes were considered statistically significant when p<0.05.

## Data availability statement

For original data, please contact.

## Acknowledgements

This project was supported by the Medical College of Wisconsin Center for Immunology (RLA) and the Neuroscience Research Center (ADE). C9ORF72 iPSCs were obtained from the Cedar Sinai Stem Cell Core. Figures 3A and graphical abstract created using BioRender.

## Author Contributions

RLA performed and analyzed experiments. All authors designed experiments and interpreted data. ADE supervised the study and provided funding. RLA wrote the manuscript and created figures. All authors edited and approved the manuscript.

## References

1. Chiò A, Logroscino G, Hardiman O, et al. Prognostic factors in ALS: A critical review. Amyotroph Lateral Scler. 2009;10(5-6):310–23. doi:10.3109/17482960802566824

2. Mejzini R, Flynn LL, Pitout IL, Fletcher S, Wilton SD, Akkari PA. ALS Genetics, Mechanisms, and Therapeutics: Where Are We Now? Front Neurosci. 2019;13:1310. doi:10.3389/fnins.2019.01310

3. Clement AM, Nguyen MD, Roberts EA, et al. Wild-type nonneuronal cells extend survival of SOD1 mutant motor neurons in ALS mice. Science. Oct 03 2003;302(5642):113–7. doi:10.1126/science.1086071

4. Birger A, Ben-Dor I, Ottolenghi M, et al. Human iPSC-derived astrocytes from ALS patients with mutated C9ORF72 show increased oxidative stress and neurotoxicity. EBioMedicine. Dec 2019;50:274–289. doi:10.1016/j.ebiom.2019.11.026

5. Reischauer C, Gutzeit A, Neuwirth C, et al. In-vivo evaluation of neuronal and glial changes in amyotrophic lateral sclerosis with diffusion tensor spectroscopy. Neuroimage Clin. 2018;20:993–1000. doi:10.1016/j.nicl.2018.10.001

6. Wong CO, Venkatachalam K. Motor neurons from ALS patients with mutations in C9ORF72 and SOD1 exhibit distinct transcriptional landscapes. Hum Mol Genet. Aug 15 2019;28(16):2799–2810. doi:10.1093/hmg/ddz104

7. Zhao C, Devlin AC, Chouhan AK, et al. Mutant C9orf72 human iPSC-derived astrocytes cause non-cell autonomous motor neuron pathophysiology. Glia. May 2020;68(5):1046–1064. doi:10.1002/glia.23761

8. Madill M, McDonagh K, Ma J, et al. Amyotrophic lateral sclerosis patient iPSC-derived astrocytes impair autophagy via non-cell autonomous mechanisms. Mol Brain. Jun 13 2017;10(1):22. doi:10.1186/s13041-017-0300-4

9. Haidet-Phillips AM, Hester ME, Miranda CJ, et al. Astrocytes from familial and sporadic ALS patients are toxic to motor neurons. Nat Biotechnol. Aug 2011;29(9):824–8. doi:10.1038/nbt.1957

10. Boillée S, Yamanaka K, Lobsiger CS, et al. Onset and progression in inherited ALS determined by motor neurons and microglia. Science. Jun 2006;312(5778):1389–92. doi:10.1126/science.1123511

11. Frakes AE, Ferraiuolo L, Haidet-Phillips AM, et al. Microglia induce motor neuron death via the classical NF-κB pathway in amyotrophic lateral sclerosis. Neuron. Mar 2014;81(5):1009–1023. doi:10.1016/j.neuron.2014.01.013

12. Guttenplan KA, Weigel MK, Adler DI, et al. Knockout of reactive astrocyte activating factors slows disease progression in an ALS mouse model. Nat Commun. 07 2020;11(1):3753. doi:10.1038/s41467-020-17514-9

13. Ziff OJ, Clarke BE, Taha DM, Crerar H, Luscombe NM, Patani R. Meta-analysis of human and mouse ALS astrocytes reveals multi-omic signatures of inflammatory reactive states. Genome Res. Jan 2022;32(1):71–84. doi:10.1101/gr.275939.121

14. McCombe PA, Henderson RD. The Role of immune and inflammatory mechanisms in ALS. Curr Mol Med. Apr 2011;11(3):246–54. doi:10.2174/156652411795243450

15. Butovsky O, Siddiqui S, Gabriely G, et al. Modulating inflammatory monocytes with a unique microRNA gene signature ameliorates murine ALS. J Clin Invest. Sep 2012;122(9):3063–87. doi:10.1172/JCI62636

16. Nagata T, Nagano I, Shiote M, et al. Elevation of MCP-1 and MCP-1/VEGF ratio in cerebrospinal fluid of amyotrophic lateral sclerosis patients. Neurol Res. Dec 2007;29(8):772–6. doi:10.1179/016164107X229795

17. Jara JH, Gautam M, Kocak N, et al. MCP1-CCR2 and neuroinflammation in the ALS motor cortex with TDP-43 pathology. J Neuroinflammation. Oct 30 2019;16(1):196. doi:10.1186/s12974-019-1589-y

18. Kuhle J, Lindberg RL, Regeniter A, et al. Increased levels of inflammatory chemokines in amyotrophic lateral sclerosis. Eur J Neurol. Jun 2009;16(6):771–4. doi:10.1111/j.1468-1331.2009.02560.x

19. Baron P, Bussini S, Cardin V, et al. Production of monocyte chemoattractant protein-1 in amyotrophic lateral sclerosis. Muscle Nerve. Oct 2005;32(4):541–4. doi:10.1002/mus.20376

20. RL A, JW A, J A, et al. Microglia Influence Neurofilament Deposition in ALS iPSC-Derived Motor Neurons. Genes. 2022;13(2):241. doi:10.3390/genes13020241

21. Gravel M, Béland LC, Soucy G, et al. IL-10 Controls Early Microglial Phenotypes and Disease Onset in ALS Caused by Misfolded Superoxide Dismutase 1. J Neurosci. Jan 20 2016;36(3):1031–48. doi:10.1523/JNEUROSCI.0854-15.2016

22. Norden DM, Fenn AM, Dugan A, Godbout JP. TGFβ produced by IL-10 redirected astrocytes attenuates microglial activation. Glia. Jun 2014;62(6):881–95. doi:10.1002/glia.22647

23. Allison RL, Welby E, Khayrullina G, Burnett BG, Ebert AD. Viral mediated knockdown of GATA6 in SMA iPSC-derived astrocytes prevents motor neuron loss and microglial activation. Glia. Jan 28 2022;doi:10.1002/glia.24153

24. Welby E, Ebert AD. Diminished motor neuron activity driven by abnormal astrocytic EAAT1 glutamate transporter activity in spinal muscular atrophy is not fully restored after lentiviral SMN delivery. Glia. May 2023;71(5):1311–1332. doi:10.1002/glia.24340

25. Franco-Bocanegra DK, McAuley C, Nicoll JAR, Boche D. Molecular Mechanisms of Microglial Motility: Changes in Ageing and Alzheimer’s Disease. Cells. 06 2019;8(6)doi:10.3390/cells8060639

26. Fu R, Shen Q, Xu P, Luo JJ, Tang Y. Phagocytosis of microglia in the central nervous system diseases. Mol Neurobiol. Jun 2014;49(3):1422–34. doi:10.1007/s12035-013-8620-6

27. Klawonn AM, Fritz M, Castany S, et al. Microglial activation elicits a negative affective state through prostaglandin-mediated modulation of striatal neurons. Immunity. Feb 2021;54(2):225–234.e6. doi:10.1016/j.immuni.2020.12.016

28. Kiskinis E, Sandoe J, Williams LA, et al. Pathways disrupted in human ALS motor neurons identified through genetic correction of mutant SOD1. Cell Stem Cell. Jun 05 2014;14(6):781–95. doi:10.1016/j.stem.2014.03.004

29. Bhinge A, Namboori SC, Zhang X, VanDongen AMJ, Stanton LW. Genetic Correction of SOD1 Mutant iPSCs Reveals ERK and JNK Activated AP1 as a Driver of Neurodegeneration in Amyotrophic Lateral Sclerosis. Stem Cell Reports. Apr 11 2017;8(4):856–869. doi:10.1016/j.stemcr.2017.02.019

30. Dafinca R, Scaber J, Ababneh N, et al. C9orf72 Hexanucleotide Expansions Are Associated with Altered Endoplasmic Reticulum Calcium Homeostasis and Stress Granule Formation in Induced Pluripotent Stem Cell-Derived Neurons from Patients with Amyotrophic Lateral Sclerosis and Frontotemporal Dementia. Stem Cells. Aug 2016;34(8):2063–78. doi:10.1002/stem.2388

31. Devlin AC, Burr K, Borooah S, et al. Human iPSC-derived motoneurons harbouring TARDBP or C9ORF72 ALS mutations are dysfunctional despite maintaining viability. Nat Commun. Jan 12 2015;6:5999. doi:10.1038/ncomms6999

32. Naujock M, Stanslowsky N, Bufler S, et al. 4-Aminopyridine Induced Activity Rescues Hypoexcitable Motor Neurons from Amyotrophic Lateral Sclerosis Patient-Derived Induced Pluripotent Stem Cells. Stem Cells. Jun 2016;34(6):1563–75. doi:10.1002/stem.2354

33. Riley JK, Takeda K, Akira S, Schreiber RD. Interleukin-10 receptor signaling through the JAK-STAT pathway. Requirement for two distinct receptor-derived signals for anti-inflammatory action. J Biol Chem. Jun 04 1999;274(23):16513–21. doi:10.1074/jbc.274.23.16513

34. Staples KJ, Bergmann M, Barnes PJ, Newton R. Stimulus-specific inhibition of IL-5 by cAMP-elevating agents and IL-10 reveals differential mechanisms of action. Biochem Biophys Res Commun. Jul 14 2000;273(3):811–5. doi:10.1006/bbrc.2000.3023

35. Abiusi E, Infante P, Cagnoli C, et al. SMA-miRs (miR-181a-5p, –324-5p, and –451a) are overexpressed in spinal muscular atrophy skeletal muscle and serum samples. Elife. 09 20 2021;10 doi:10.7554/eLife.68054

36. Lobo-Silva D, Carriche GM, Castro AG, Roque S, Saraiva M. Balancing the immune response in the brain: IL-10 and its regulation. J Neuroinflammation. 11 24 2016;13(1):297. doi:10.1186/s12974-016-0763-8

37. Zhou Z, Peng X, Insolera R, Fink DJ, Mata M. Interleukin-10 provides direct trophic support to neurons. J Neurochem. Sep 2009;110(5):1617–27. doi:10.1111/j.1471-4159.2009.06263.x

38. Spooren A, Kolmus K, Laureys G, et al. Interleukin-6, a mental cytokine. Brain Res Rev. Jun 24 2011;67(1-2):157–83. doi:10.1016/j.brainresrev.2011.01.002

39. Morrice JR, Gregory-Evans CY, Shaw CA. Animal models of amyotrophic lateral sclerosis: A comparison of model validity. Neural Regen Res. Dec 2018;13(12):2050–2054. doi:10.4103/1673-5374.241445

40. Ferraiuolo L, Maragakis NJ. Mini-Review: Induced pluripotent stem cells and the search for new cell-specific ALS therapeutic targets. Neurosci Lett. Jun 11 2021;755:135911. doi:10.1016/j.neulet.2021.135911

41. Ragagnin AMG, Shadfar S, Vidal M, Jamali MS, Atkin JD. Motor Neuron Susceptibility in ALS/FTD. Front Neurosci. 2019;13:532. doi:10.3389/fnins.2019.00532

42. Izrael M, Slutsky SG, Revel M. Rising Stars: Astrocytes as a Therapeutic Target for ALS Disease. Front Neurosci. 2020;14:824. doi:10.3389/fnins.2020.00824

43. Ng W, Ng SY. Remodeling of astrocyte secretome in amyotrophic lateral sclerosis: uncovering novel targets to combat astrocyte-mediated toxicity. Transl Neurodegener. Dec 26 2022;11(1):54. doi:10.1186/s40035-022-00332-y

44. Philips T, Rothstein JD. Glial cells in amyotrophic lateral sclerosis. Exp Neurol. Dec 2014;262 Pt B:111–20. doi:10.1016/j.expneurol.2014.05.015

45. Filipi T, Hermanova Z, Tureckova J, Vanatko O, Anderova AM. Glial Cells-The Strategic Targets in Amyotrophic Lateral Sclerosis Treatment. J Clin Med. Jan 18 2020;9(1) doi:10.3390/jcm9010261

46. Cassina P, Miquel E, Martínez-Palma L, Cassina A. Glial Metabolic Reprogramming in Amyotrophic Lateral Sclerosis. Neuroimmunomodulation. 2021;28(4):204–212. doi:10.1159/000516926

47. Conductier G, Blondeau N, Guyon A, Nahon JL, Rovère C. The role of monocyte chemoattractant protein MCP1/CCL2 in neuroinflammatory diseases. J Neuroimmunol. Jul 27 2010;224(1-2):93–100. doi:10.1016/j.jneuroim.2010.05.010

48. Strickland MR, Ibanez KR, Yaroshenko M, Diaz CC, Borchelt DR, Chakrabarty P. IL-10 based immunomodulation initiated at birth extends lifespan in a familial mouse model of amyotrophic lateral sclerosis. Sci Rep. Nov 30 2020;10(1):20862. doi:10.1038/s41598-020-77564-3

49. Lorenzini I, Alsop E, Levy J, et al. Moderate intrinsic phenotypic alterations in. Front Cell Neurosci. 2023;17:1179796. doi:10.3389/fncel.2023.1179796

50. Smeyers J, Banchi EG, Latouche M. C9ORF72: What It Is, What It Does, and Why It Matters. Front Cell Neurosci. 2021;15:661447. doi:10.3389/fncel.2021.661447

51. Rostalski H, Leskelä S, Huber N, et al. Astrocytes and Microglia as Potential Contributors to the Pathogenesis of. Front Neurosci. 2019;13:486. doi:10.3389/fnins.2019.00486

52. Taha DM, Clarke BE, Hall CE, et al. Astrocytes display cell autonomous and diverse early reactive states in familial amyotrophic lateral sclerosis. Brain. Apr 18 2022;145(2):481–489. doi:10.1093/brain/awab328

53. Dimayuga FO, Wang C, Clark JM, Dimayuga ER, Dimayuga VM, Bruce-Keller AJ. SOD1 overexpression alters ROS production and reduces neurotoxic inflammatory signaling in microglial cells. J Neuroimmunol. Jan 2007;182(1-2):89–99. doi:10.1016/j.jneuroim.2006.10.003

54. Simpson DSA, Oliver PL. ROS Generation in Microglia: Understanding Oxidative Stress and Inflammation in Neurodegenerative Disease. Antioxidants (Basel). Aug 13 2020;9(8) doi:10.3390/antiox9080743

55. Panov A, Dikalov S, Shalbuyeva N, Hemendinger R, Greenamyre JT, Rosenfeld J. Species– and tissue-specific relationships between mitochondrial permeability transition and generation of ROS in brain and liver mitochondria of rats and mice. Am J Physiol Cell Physiol. Feb 2007;292(2):C708–18. doi:10.1152/ajpcell.00202.2006

56. Caso JR, Pradillo JM, Hurtado O, Lorenzo P, Moro MA, Lizasoain I. Toll-like receptor 4 is involved in brain damage and inflammation after experimental stroke. Circulation. Mar 27 2007;115(12):1599–608. doi:10.1161/CIRCULATIONAHA.106.603431

57. Spiller KJ, Restrepo CR, Khan T, et al. Microglia-mediated recovery from ALS-relevant motor neuron degeneration in a mouse model of TDP-43 proteinopathy. Nat Neurosci. 03 2018;21(3):329–340. doi:10.1038/s41593-018-0083-7

58. Galbiati M, Crippa V, Rusmini P, et al. Multiple Roles of Transforming Growth Factor Beta in Amyotrophic Lateral Sclerosis. Int J Mol Sci. Jun 16 2020;21(12)doi:10.3390/ijms21124291

59. Bae JS, Simon NG, Menon P, Vucic S, Kiernan MC. The puzzling case of hyperexcitability in amyotrophic lateral sclerosis. J Clin Neurol. Apr 2013;9(2):65–74. doi:10.3988/jcn.2013.9.2.65

60. Menon P, Higashihara M, van den Bos M, Geevasinga N, Kiernan MC, Vucic S. Cortical hyperexcitability evolves with disease progression in ALS. Ann Clin Transl Neurol. May 2020;7(5):733–741. doi:10.1002/acn3.51039

61. Seminary ER, Sison SL, Ebert AD. Modeling Protein Aggregation and the Heat Shock Response in ALS iPSC-Derived Motor Neurons. Front Neurosci. 2018;12:86. doi:10.3389/fnins.2018.00086

62. Seminary ER, Santarriaga S, Wheeler L, et al. Motor Neuron Generation from iPSCs from Identical Twins Discordant for Amyotrophic Lateral Sclerosis. Cells. 02 2020;9(3)doi:10.3390/cells9030571

63. Luecke IW, Lin G, Santarriaga S, Scaglione KM, Ebert AD. Viral vector gene delivery of the novel chaperone protein SRCP1 to modify insoluble protein in in vitro and in vivo models of ALS. Gene Ther. Jul 08 2021; doi:10.1038/s41434-021-00276-4

64. McQuade A, Coburn M, Tu CH, Hasselmann J, Davtyan H, Blurton-Jones M. Development and validation of a simplified method to generate human microglia from pluripotent stem cells. Mol Neurodegener. 12 2018;13(1):67. doi:10.1186/s13024-018-0297-x

65. Maury Y, Côme J, Piskorowski RA, et al. Combinatorial analysis of developmental cues efficiently converts human pluripotent stem cells into multiple neuronal subtypes. Nat Biotechnol. Jan 2015;33(1):89–96. doi:10.1038/nbt.3049

66. Allison R, Suneja M, LaCroix M, Harmelink M, Ebert A. IL-1Ra and CCL5, but not IL-10, are promising targets for treating SMA astrocyte-driven pathology. BioRxiv 2023.

