## Supplemental tables and figures for "ALS iPSC-derived microglia and motor neurons respond to astrocyte-targeted IL-10 and CCL2 modulation"

Supplementary Materials – Astrocyte-targeted therapeutics for ALS

### Supplemental Table S1 – Primer sequences

| Target Gene | Forward Sequence | Reverse Sequence |
|---|---|---|
| GFAP | GTCCCCCACCTAGTTTGCAG | TAGTCGTTGGCTTCGTGCTT |
| ALDH1L1 | GCCTGGCTTCTGGTGTCTTC | GCCACGTCGGTCTTGTTGTA |
| NFkB | GCAGCACTACTTCTTGACCACC | TCTGCTCCTGAGCATTGACGTC |
| IL-1B | CCACAGACCTTCCAGGAGAATG | GTGCAGTTCAGTGATCGTACAGG |
| IL-6 | AGACAGCCACTCACCTCTTCAG | TTCTGCCAGTGCCTCTTTGCTG |
| C1q | CAACACAGGCTGCTACGGGATC | CTGCCCTTTGGGTCCTCGGAT |
| C3 | GTGGAAATCCGAGCCGTTCTCT | GATGGTTACGGTCTGCTGGTGA |
| GDNF | CGCCGAAGACCGCTCCCTCG | ATCCATGACATCATCGAACTGATC |
| BDNF | CATCCGAGGACAAGGTGGCTTG | GCCGAACTTTCTGGTCCTCATC |

Supplemental Figure S1 – Additional qRT-PCR targets in SOD1 and C9ORF72 astrocytes after IL10CCL2 treatment.

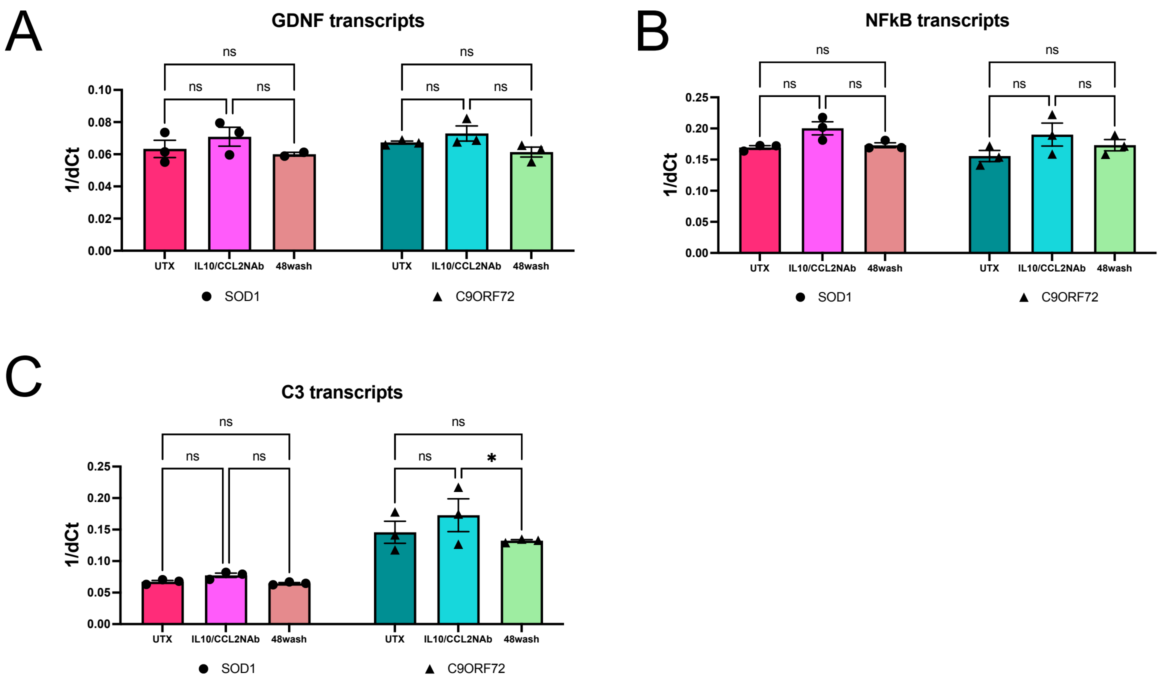


**Supplemental Figure S1 – Additional qRT-PCR targets in SOD1 and C9ORF72 astrocytes after IL10/CCL2NAb treatment.** IL10/CCL2NAb treated SOD1 and C9ORF72 astrocytes do not have altered expression of transcripts for GDNF (**A**) or NFkB (**B**). **C**. C9ORF72 astrocytes do have decreased transcripts for C3 48-hours after IL10CCL2 treatment whereas SOD1 astrocytes do not. 2-way ANOVAs, *$p<0.05$.

Supplemental Figure S2 – Addition of microglia to untreated MN cultures does not alter calcium flux.

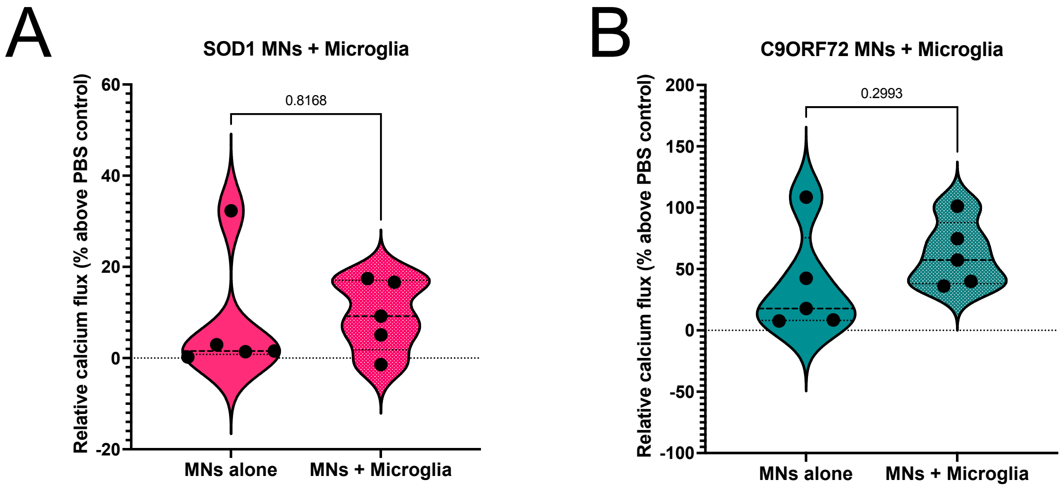


**Supplemental Figure S2 – Addition of microglia to untreated MN cultures does not alter calcium flux.** No significant differences in calcium flux in SOD1 (**A**) or C9ORF72 (**B**) motor neurons after addition of microglia without astrocyte influence. T-tests, ns.

Supplemental Figure S3 – TUNEL only channel image exports of representative images.

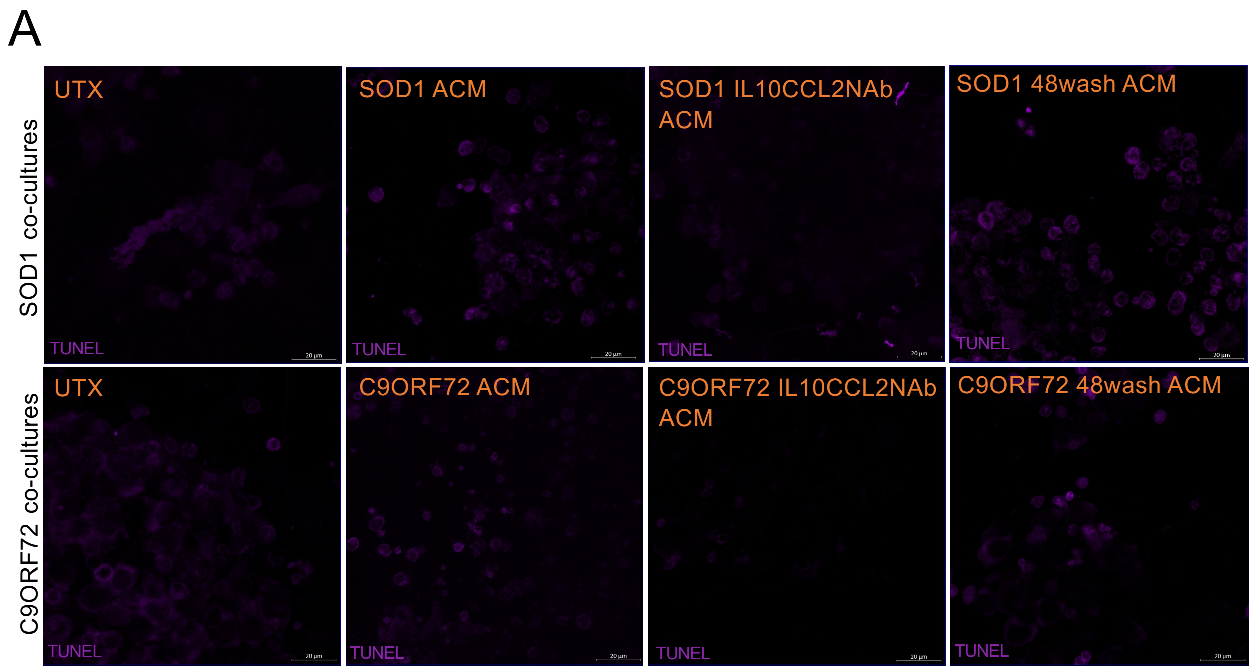


**Supplemental Figure S3 – TUNEL only channel image exports of representative images. A**. TUNEL channel only exports from Figure 5E representative images taken with Zeiss confocal microscope using a 63x oil objective.
